# Adult Human Heart ECM Improves Human iPSC-CM Function via Mitochondrial and Metabolic Maturation

**DOI:** 10.1101/2023.10.31.565062

**Authors:** S. Gulberk Ozcebe, Mateo Tristan, Pinar Zorlutuna

## Abstract

Myocardial infarction can lead to the loss of billions of cardiomyocytes, and while cell-based therapies are a promising option, the immature nature of in vitro-generated human induced pluripotent stem cell (iPSC)-derived cardiomyocytes (iCMs) is a significant roadblock to their development. Through the years, various approaches have emerged to improve iCM maturation, yet none could fully recapitulate the complexity of cardiac development and were not enough to achieve full cardiac maturity *in vitro*.

Cardiac differentiation occurs at the early stages of development in a highly dynamic environment. Although significantly improved over the past two decades, small molecule-based iPSC differentiation protocols don’t go beyond producing high purity fetal iCMs. Recently adult extracellular matrix (ECM) was shown to retain tissue memory and has shown some success in driving tissue-specific differentiation in unspecified cells in various organ systems. Therefore, here, we first characterized the adult human heart left ventricle components. We then investigated the effect of adult human heart-derived ECM on iPSC cardiac differentiation and subsequent maturation. By preconditioning iPSCs with ECM, we tested whether creating a cardiac environment around iPSCs would drive them toward cardiac fate before small molecule mediated differentiation. Ultimately, we investigated ECM components that might be responsible for the observed effects.

We identified critical glycoproteins and proteoglycans involved in early cardiac development in the adult heart ECM. Namely, adult ECM had extracellular galactin-1, fibronectin, fibrillins, and basement-membrane-specific heparan-sulfate proteoglycan (HSPG), which have been implicated in normal heart development and associated with various embryonic developmental processes.

Relatedly, we showed that preconditioning iPSCs with adult ECM resulted in enhanced cardiac differentiation, yielding iCMs with higher functional maturity. Further investigation revealed that a more developed mitochondrial network and coverage as well as enhanced metabolic maturity and a shift towards a more energetic profile allowed the observed functional enhancement in ECM pretreated iCMs.

These findings demonstrate the potential of using cardiac ECM for promoting iCM maturation and suggest a promising strategy for improving the development of iCM-based therapies and in vitro cardiac disease modeling and drug screening studies. Upon manipulating ECM, such as heat denaturation and sonication to eliminate protein components and release ECM bound vesicle contents, respectively, we concluded that the beneficial effects that we observed are not solely due to the ECM proteins, and might be related to the decorative units attached to them.

## 1. Introduction

Myocardial infarction causes the loss of up to 1 billion cardiomyocytes within a few hours[1]. Since myocardium has negligibly low regenerative abilities, cell-based therapies are offered to repopulate the lost cell body in the infarct region. Recently human induced pluripotent stem cell (hiPSC)-derived cardiomyocyte (iCM) therapies have been an attractive approach. However, iCMs generated *in vitro* remain immature, resembling fetal rather than adult hearts with both physical and functional properties[2,3]. Therefore, iCM immaturity remains the biggest roadblock in developing iCM-based therapies, as well as *in vitro* cardiac disease modeling and drug screening studies.

Various approaches have been proposed to promote the maturation of iCM cultures[4–9]. As one of the early strategies, prolonged culture of iCMs for up to 120 days led to structural changes such as increased cell size and increased myofibrillar density and alignment, as well as improvements in contractile performance, calcium handling, and electrophysiological properties[2,10,11]. However, it has been demonstrated that cardiac maturity is not directly proportional to the duration of culture time. Despite their structural maturation, an extended culture of iCMs beyond 3 months leads to reduced cardiac function and increased expression of aging and stress-related markers [3]. Additionally, the maintenance of cells for prolonged periods in culture is time-consuming and costly, making it a less favorable approach.

The heart is one of the first organs to form and function during embryogenesis[12]. Cardiac differentiation occurs at the early stages of development, where transcription factors drive the mesodermal to cardiac progenitor, and cardiac progenitor to cardiomyocyte differentiation, along with non-beating cardiac cells. CMs undergo structural, electrophysiological, transcriptional and metabolic changes during the maturation process. While it can take months in rodents, CM maturation takes around 6 years in humans to complete [13]. It is therefore extremely challenging to recapitulate human CM maturation *in vitro.* Small molecule-based iPSC differentiation protocols have improved over the past two decades for reproducible high purity cardiac populations, However, using extracellular cues to mimic the cardiac environment and enhance iCM maturation is a new approach.

Decellularized extracellular matrix (ECM) from various organs, including the heart, was shown to contain tissue-specific cues that play a crucial role in regulating homeostasis, cell response, and development[3,14]. Recellularization of three-dimensional whole organ scaffolds with iPSCs has been explored on the same premises that ECM memorizes its tissue origin and provides cues that derive organotypic differentiation processes. Decellularized three different human organs including heart ECM slices were reported to direct organ-specific differentiation of uncommitted iPSC-derived mesoderm cells[15]. Similarly, powdered decellularized human atrial and ventricular ECMs were reported to induce heart chamber-specific cardiac differentiation of human iPSCs [16]. Researchers further suggested that ventricular ECM may promote electrical maturation of the ventricular CMs, however, this has not been investigated further.

We previously showed the bioactive nature of cardiac ECM and the dynamic compositional shifts observed with the ECM age[2]. The results from human and mouse studies[15,17,18] are in alignment with our previous study reporting that the ‘adult ECM’ contains biochemical memory that promotes tissue-specific differentiation. Therefore, here, we’ve investigated the effect of adult human heart-derived ECM on iPSC differentiation and subsequent cardiac maturation by driving iPSCs toward cardiac fate before the small molecule mediated differentiation. We treated iPSCs with adult heart ECM for 5 days before the differentiation and followed the differentiation protocol with no modifications. On day 30, we investigated the iCM maturity at both cellular and subcellular levels, including transcriptional changes, contractile function, mitochondrial structure, and energy metabolism. We showed that adult cardiac ECM preconditioning enhanced the iPSC cardiac differentiation *in vitro* in a dose dependent manner, and yielded iCMs that expressed transcription factors related to a more mature phenotype both functionally and structurally. Additionally, ECM preconditioned iCMs displayed a more energetic metabolic state, supported by increased mitochondrial coverage, interconnected mitochondrial network structure and faster beating. Establishing the effects of ECM preconditioning on iPSCs, we then investigated potential ECM components to be responsible for the observed effects. ECM was denatured and sonicated to test the protein and ECM-bound extracellular vesicle (EV) components of ECM, respectively. As denaturing ECM did not nullify the ECM pretreatment effects, our results suggest that ECM proteins are not the only factor driving the observed effects.

## 2. Materials and Methods

### 2.1. Human heart collection

De-identified human hearts that were deemed unsuitable for transplantation and donated to research, were acquired from Indiana Donor Network under the Institutional Review Board (IRB) approval for deceased donor tissue recovery. Human heart tissues were grouped as adult (from 30-50-years-old patients, n=3). For storage, hearts were dissected into their chambers and kept separately in a −80°C freezer until use. We only used the young left ventricles (n=3) for the ECM pretreatments.

### 2.2. Decellularization of human heart tissue for matrix preparation

Left ventricles from adult donors were sectioned and decellularized following the previous decellularization protocol [19]. Briefly, we first stripped the fatty tissue around the left ventricular myocardial tissue and sliced the tissues into thin sections (<1mm). To decellularize, tissues were washed in 1% (wt/vol) sodium dodecyl sulfate (SDS) (VWR, #97062) for 24 hours or until white transparent tissue was obtained, then in 1% (wt/vol) Triton 100-X (Sigma-Aldrich, #A16046) for 30 minutes. After decellularization, samples were washed thoroughly with DI water to remove any residual detergent. To delipidize, tissues were washed with the isopropanol (IPA) for 3 hours then rehydrated in DI and treated with 50U/ml DNase (Millipore Sigma, #10104159001) for 8 hours followed by an overnight DI rinse. All steps were conducted with constant agitation at RT.

Prepared ECMs were lyophilized and pulverized with liquid nitrogen. ECM powder was digested in 1 mg/mL pepsin (Sigma-Aldrich, #P6887) in 0.1M HCl (10:1, w/w, dry ECM:pepsin) at RT with constant stirring until a homogeneous solution was obtained. The insoluble remnants were removed by centrifugation, the supernatant was neutralized using 1M NaOH solution, and used immediately to prevent degradation. Prior to experiments, we measured the total protein concentrations using Rapid Gold BCA Assay (Thermo Scientific, # A53227) and diluted ECM solutions to either 0.01 mg/ml (1x) or 0.05 mg/ml (5x) with the culture media.

### 2.3. Alternative decellularization and solubilization protocol for EV preservation

Decellularization and digestion were performed in accordance with current standards for maintaining EV integrity[20]. Tissues were washed in 0.1% peracetic acid (Sigma Aldrich, USA) and 4% ethanol (Sigma Aldrich) at 200 rpm for 2 h, then rinsed in phosphate buffered saline (PBS) at 200 rpm for 2 h, and then washed again in peracetic acid/ethanol solution at 200 rpm for 16 h. Decellularized ECMs were rinsed with PBS and then with DI water. Prepared ECMs were lyophilized and pulverized with liquid nitrogen. ECM powder was digested in digestion buffer containing 0.1 mg/mL collagenase type II (Corning), 50 mM Tris buffer (Sigma Aldrich), 5 mM CaCl2 (Amresco), and 200 mM of NaCl (Sigma Aldrich) at RT with constant stirring for 24h or until few or no solid particles could be observed in the solution, with brief remixing every 6-8 h.

### 2.4. Vesicle Extraction and Isolation

Digested ECM was centrifuged three times at 500g for 10 min, 2500g for 20 min, and 10,000g for 30 min, and the pellet was discarded after each centrifugation step to remove any remaining insoluble matrix remnants. The final supernatant was centrifuged at 100,000g at 4°C for 70 min using an ultracentrifuge (Optima MAX-XP Tabletop Ultracentrifuge, Beckman Coulter). The pellet was either used immediately or stored dry at −80°C. A final concentration of 12.5 ug/ml is used to treat iPSCs[21].

### 2.5. ECM manipulations

ECM was heat denatured at 80C for 3 hours in a wet environment as suggested by literature [22]. ECM was sonicated using an ultrasonicator at the following settings: 30% amplitude, 20 sec total time, 0.5 sec ON/1 sec OFF pulses, over ice.

### 2.6. Mass spectroscopy

Mass spectrometry (MS) was used to determine the protein composition of the adult human heart left ventricles (n=3). Tissues were digested in the digest solution (5M urea, 2M thiourea, 100mM Ammonium bicarbonate and 50mM Dithiothreitol, pH 8.0) at 4°C for 24 h with constant stirring. The resulting soluble ECM components were processed via in-solution trypsin digestion [23]. The protein concentration of the samples was measured with the Pierce Rapid Gold BCA Assay (Thermo Scientific) and equalized prior to MS. Samples were further processed using S-TrapTM mini spin column digestion protocol and eluted peptides were used for MS analysis. The ECM components for human LV were identified and quantified using Thermo Scientific Proteome Discoverer and PEAKSOnline Proteomics Server software.

### 2.7. Human iPSC cell culture

The cell line used in this study is DiPS 1016 SevA (RRID: CVCL_UK18) from human dermal fibroblasts obtained from Harvard Stem Cell Institute iPS Core Facility. Cells were cultured in humidified incubators at 37 °C and 5% CO2. Human iPS cells were cultured routinely in mTeSR-1 media (StemCell Technologies, #05825) on 1% Geltrex-coated plates (Invitrogen, #A1413201). At 80-85% confluency, cells were passaged using Accutase (StemCell Technologies, #07920) and seeded at low density 45 × 10^4^ cells/cm^2^ on well plates with Y-27632 (ROCK inhibitor, 5 *μ*M), (StemCell Technologies, #129830-38-2) in mTeSR-1 media. The culture was maintained or supplemented with ECM/EV with daily media changes until 90% confluency was reached.

### 2.8. iPSC cardiac differentiation and cell culture

Once 90% confluency was reached, cardiac differentiation was initiated following the canonical Wnt pathway[2]. To direct cardiac differentiation, cells are sequentially treated with CHIR99021 (12 *μ*M) (Stemcell Technologies, #72052) for 24 hours followed by RPMI 1640 medium with B-27 supplement without insulin (2%) (Gibco, #A1895601) (CM(-)). Cells were then treated with Wnt pathway inhibitor IWP-4 (5 µM) (Stemcell Technologies, #72552) for 48 hours followed by CM(-) for 48 hours. From day 9 on, cells were maintained in RPMI 1640 medium with B-27 (2%) (Gibco, #17504044) (CM(+)) and media was changed every 3 days.

### 2.9. Beating and calcium flux

Block-matching algorithm was performed using a custom MATLAB code. Briefly, the spontaneous beating of the iCMs was recorded as bright field videos (n=3-4 ROI per sample) and exported as a series of single-frame image files. The images were then divided into square blocks of NxN pixels. The movement of a given block at the kth frame is tracked by matching its intensity to the (k + 1)th frame within a square window of a fixed width flanking from each side. The pixel values were determined based on the speed of calculation and accuracy of the block-matching method. The matching criterion used for block movement is Mean Absolute Difference as described previously[24].

The calcium (Ca^2+^) transient of iCMs was assessed to further investigate electrophysiology. Cells in culture were washed with PBS and the media was replaced by Ca^2+^-sensitive Fluo-4 AM (Life Technologies) solution, as instructed by the manufacturer. Cell beating was recorded in real-time using a fluorescence microscope (Axio Observer.Z1, Zeiss, Hamatsu C11440 digital camera) at 30 ms exposure for 30 sec. To assess the drug response, we treated cells with isoproterenol (1 µM, ISO) for 10 min at 37°C. Spontaneous beating before and after ISO was recorded. The baseline for calcium transient and an initial beat rate of the cells were obtained from pre-drug recordings (5 images/well, and 3 wells/age group). The beating rate after ISO treatment relative to that before treatment was calculated by dividing post-ISO beating by pre-ISO beating. The rate of Ca^2+^ release (time to peak Ca^2+^ transient amplitude) and action potential duration at 50% (APD_50_) and 90% (APD_90_) of the amplitude were calculated from the raw data obtained from intensity versus time plots.

### 2.10. Mitochondrial network analysis

Mitochondria were labeled with MitoTracker following the manufacturer’s protocol. Fluorescence microscopy images of iCMs were taken with a Zeiss LSM800 laser-scanning confocal microscope and a 1.40-NA 63× objective. The mitochondrial network structure was visualized and quantified using the Mitochondria Analyzer plugin in ImageJ[25]. Mitochondria images were superimposed with the mitochondria skeleton. Skeleton images of the binary images after thresholding and noise removal were generated. With AnalyzeSkeleton plugin average mitochondrial area in a cell, the number of branches of each mitochondrion, the number of branch junctions of each mitochondrion, and the average length of branches of each mitochondrion per branch were quantified.

### 2.11. Seahorse energy profiling

The Seahorse XF96 extracellular flux analyzer was used to assess mitochondria function. The plates were coated with Matrigel and the iCMs were seeded with a density of 100,000 per XF96 well. The culture medium was exchanged for Agilent Seahorse XF RPMI Basal Medium supplemented with 2 mM glutamine, 10 mM glucose and 1 mM Sodium Pyruvate 1 hour before the assay and for the duration of the measurement. For the Mito Stress test, oligomycin (2.5 mM), FCCP (2 mM), rotenone and antimycin A (2.5 mM); for the Glycolysis stress test, glucose (10mM), oligomycin (1 mM), and 2-DG (50mM) were injected in the written order. The oxygen consumption rate (OCR) and extracellular acidification rate (ECAR) values were normalized to the number of cells captured by one 10x field of view per well quantified by Hoechst staining. The baseline OCR was defined as the average values measured from time points 1 to 3 (the last measurement before the first inhibitor injection) during the experiments. MitoATP Production Rate and glycoATP Production Rate were calculated according to Equations 1-9 from OCR and ECAR measurements, obtained using an Agilent Seahorse XFe96 Analyzer.

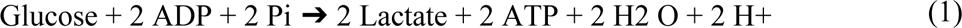

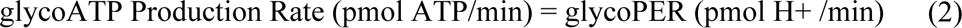

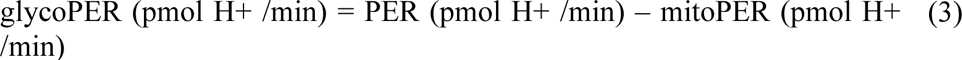

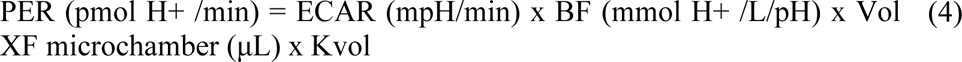

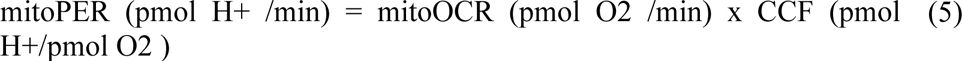

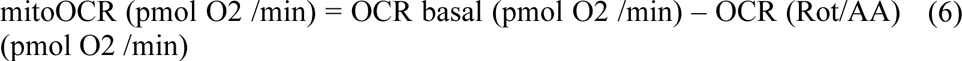

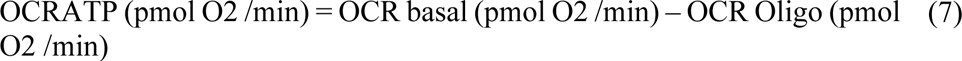

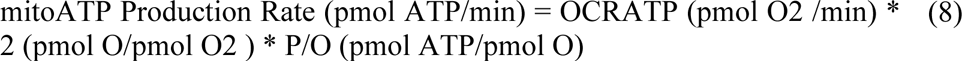

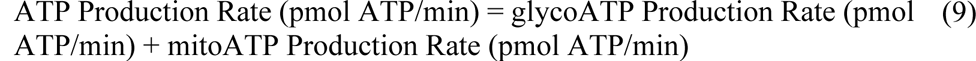

### 2.12. RT-qPCR

RNA was isolated using the RNAeasy Plus Mini Kit (Qiagen, Germany). Total RNA was isolated using the RNAeasy Plus Mini Kit (Qiagen, Germany) as per the manufacturer’s instructions. After RNA isolation the respective complementary DNA (cDNA) was then synthesized using the Bio-Rad iScript cDNA synthesis kit (Bio-Rad, USA) as per the manufacturer’s instructions. Gene expression of myosin heavy chain 6 (*MHC6*), myosin heavy chain 7 (*MHC7*), homeobox Nkx2-5 (*NKX2*), cardiac muscle troponin T (*TNNT2*), and ATPase Sarcoplasmic/Endoplasmic Reticulum Ca2+ Transporting 2 (*ATP2A2)* were quantified from 1.5 *μ*l of template cDNA using Bio-Rad SYBR Green as per manufacturer’s instructions. All assays were performed in triplicate on a CFX Connect Real-Time PCR System (Bio-Rad). Analysis was performed using the CFX Manager Software (Bio-Rad). Gene expression levels were calculated using the ΔC_T_ and normalized to GAPDH (all primers were purchased from Bio-Rad) as a housekeeping gene. For the Nanostring assay (Nanostring, USA), RNA was isolated as previously described and the assay was performed following the manufacturer’s instructions with all analysis performed using the nSolver software provided by Nanostring.

### 2.13. Statistical Analysis

The mean ± standard deviation (SD) was reported for all replicates. One-way ANOVA with post-hoc Tukey’s test was used to assess the statistically significant differences. All p values are two-sided and statistical significance was defined as p < 0.05 and sample size was (n) ≥ 3 for all experiments.

## 3. Results

### 3.1. Adult Human Heart preparation and characterization

Adult human left ventricles (LV, 30-50 years old, n=3) were sectioned and decellularized (Fig. 1A-B). To ensure human cardiac tissue decellularization with the preservation of ECM bioactivity, previous decellularization methods were optimized and a combinational protocol using ionic and non-ionic detergent washes was followed [26]. Due to the fatty nature of the human heart, an additional alcohol-based delipidation step was added following decellularization. We confirmed the absence of DNA and preservation of ECM proteins via hematoxylin and eosin (H&E), and Masson’s trichrome staining, respectively (Fig. 1C). In addition, we measured the double-stranded DNA content and verified it to be less than 50 ng/µl to ensure complete decellularization (Fig. 1D). Adult human ECM was then solubilized and the most abundant 12 ECM components are identified with mass spectroscopy (Fig. 1E). In addition to collagens, we identified critical glycoproteins and proteoglycans involved in early cardiac development (Fig. 1E and Supp. Fig. 1). Notably, extracellular galectin-1 was previously shown to induce skeletal muscle differentiation in human fetal mesenchymal stem cells [27], suggesting a potential analogous role in cardiac muscle differentiation. Similarly, fibronectin, fibrillins, and basement-membrane-specific heparan-sulfate proteoglycan (HSPG), Perlecan have been implicated in normal heart development and associated with various embryonic developmental processes[28].

**Figure 1.**
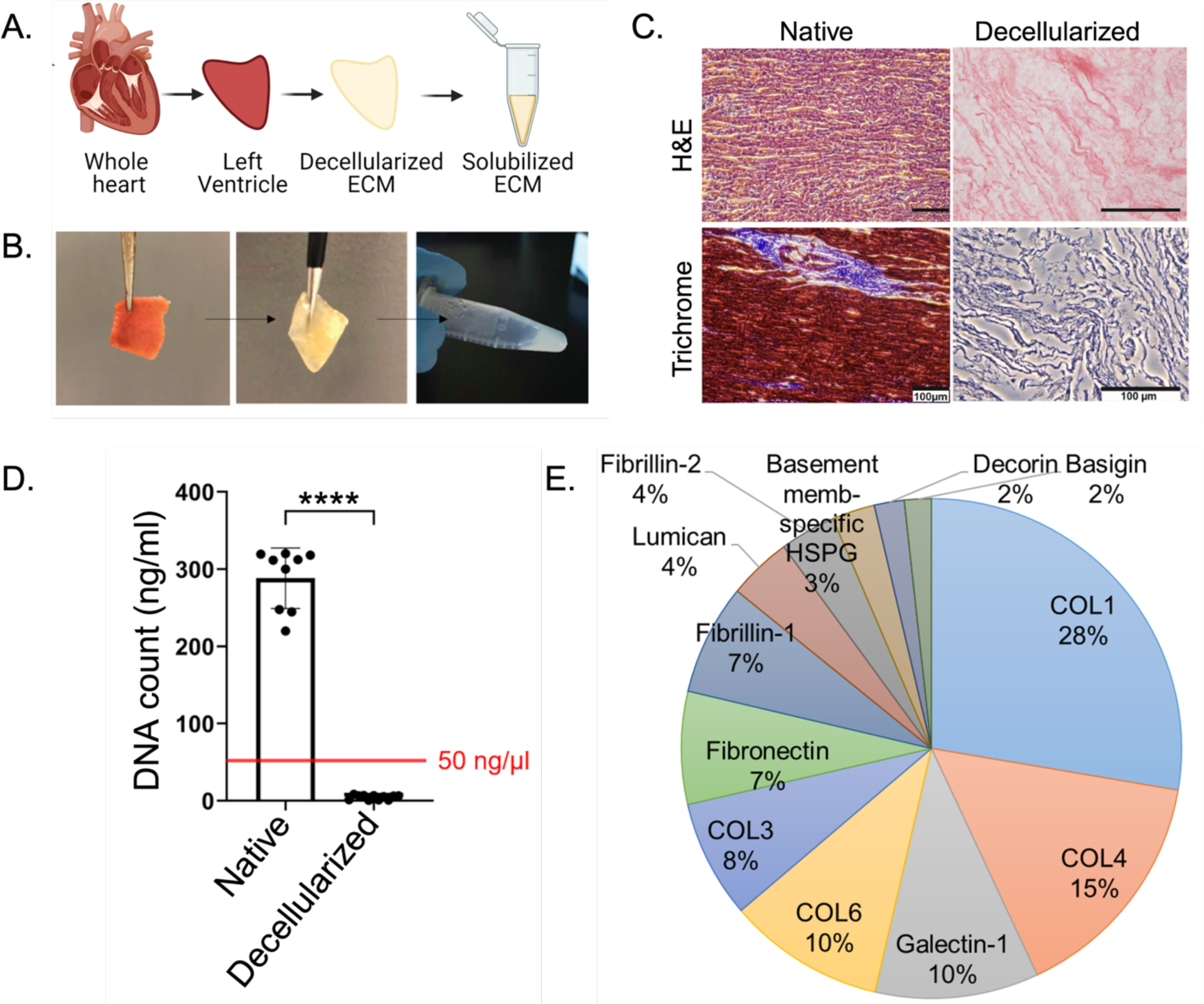
Decellularization of the native tissues and their biochemical analysis. (A) Schematic and (B) representative images demonstrating the workflow of the decellularization and solubilization process. Created by Biorender.com. (C) Hematoxylin and eosin (H&E) and Masson’s Trichrome staining, (D) Quantification of total DNA content of native and decellularized cardiac tissue. (E) 12 most abundant proteins detected by mass spectroscopy.

**Supplementary Figure 1.**
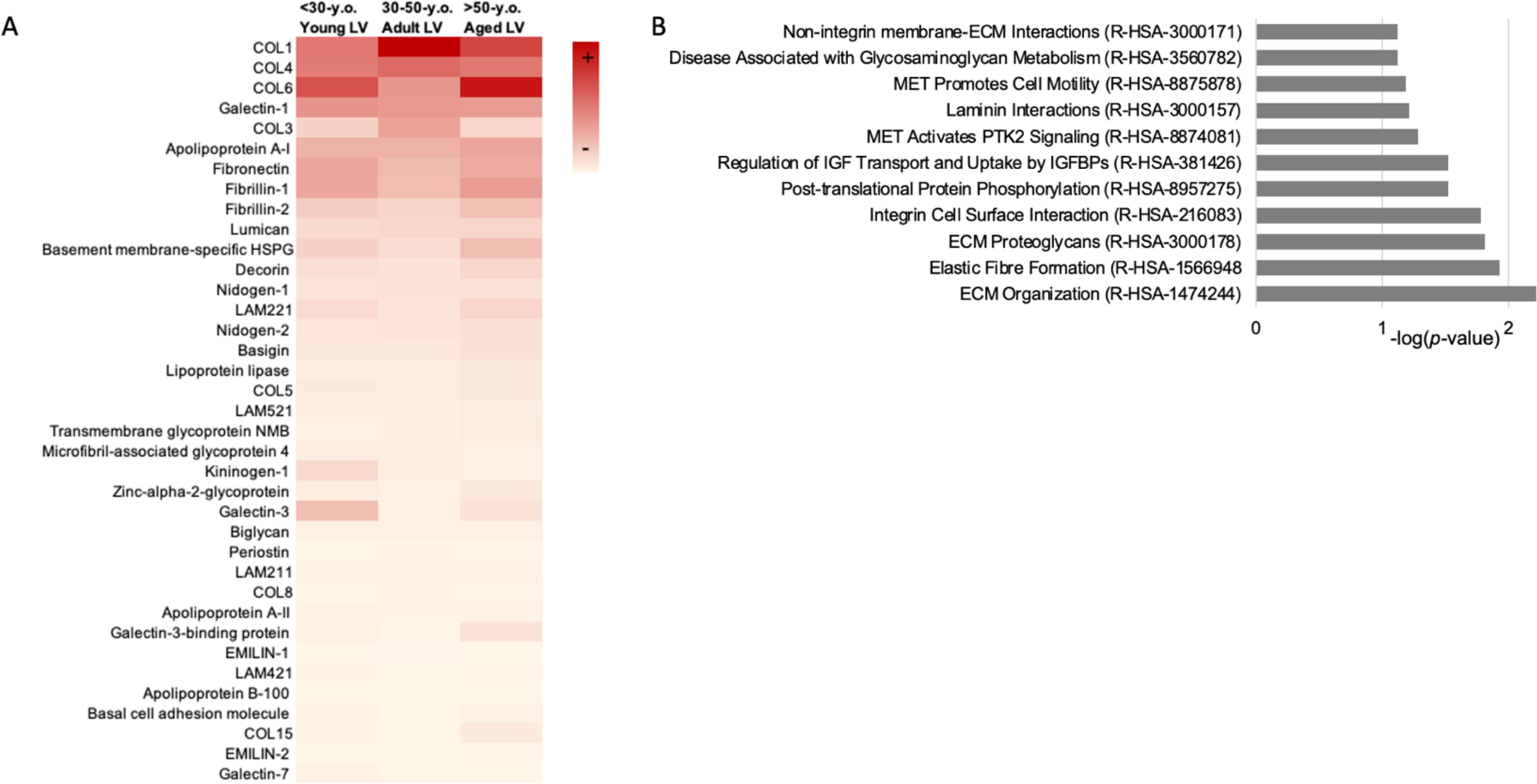
(A) Heatmap showing cardiac development associated ECM components listed with respect to adult ECM expressions in descending order. Values were calculated as the area under the peak using PEAKSOnline Proteomics Server software. (B) Top biological pathways associated with detected ECM components by Reactome Pathway Database 2022.

### 3.2. Adult ECM pretreatment improves cardiac function

We recorded the spontaneous beating of the cells and visualized the beating kinetics using vector heat maps. The maps showed that high dose ECM pretreated cells displayed tissue like beating (Fig. 2A). Although the control and pretreated groups had similar average beating areas, ECM pretreated cells displayed significantly higher beat frequency and velocity (Fig. 2B-D). We recorded the calcium transient during spontaneous beating and calculated maximum contraction (time-to-peak, TTP), 50% decay (APD50), and 90% decay durations (APD90). The high dose ECM pretreated cells exhibited a distinct ventricular-type action potential (AP) profile, characterized by a notable shoulder or late plateau (Fig. 2E). This was confirmed by the significantly prolonged APD90 in the high dose ECM pretreated cells (Fig. 2G-H).

**Figure 2.**
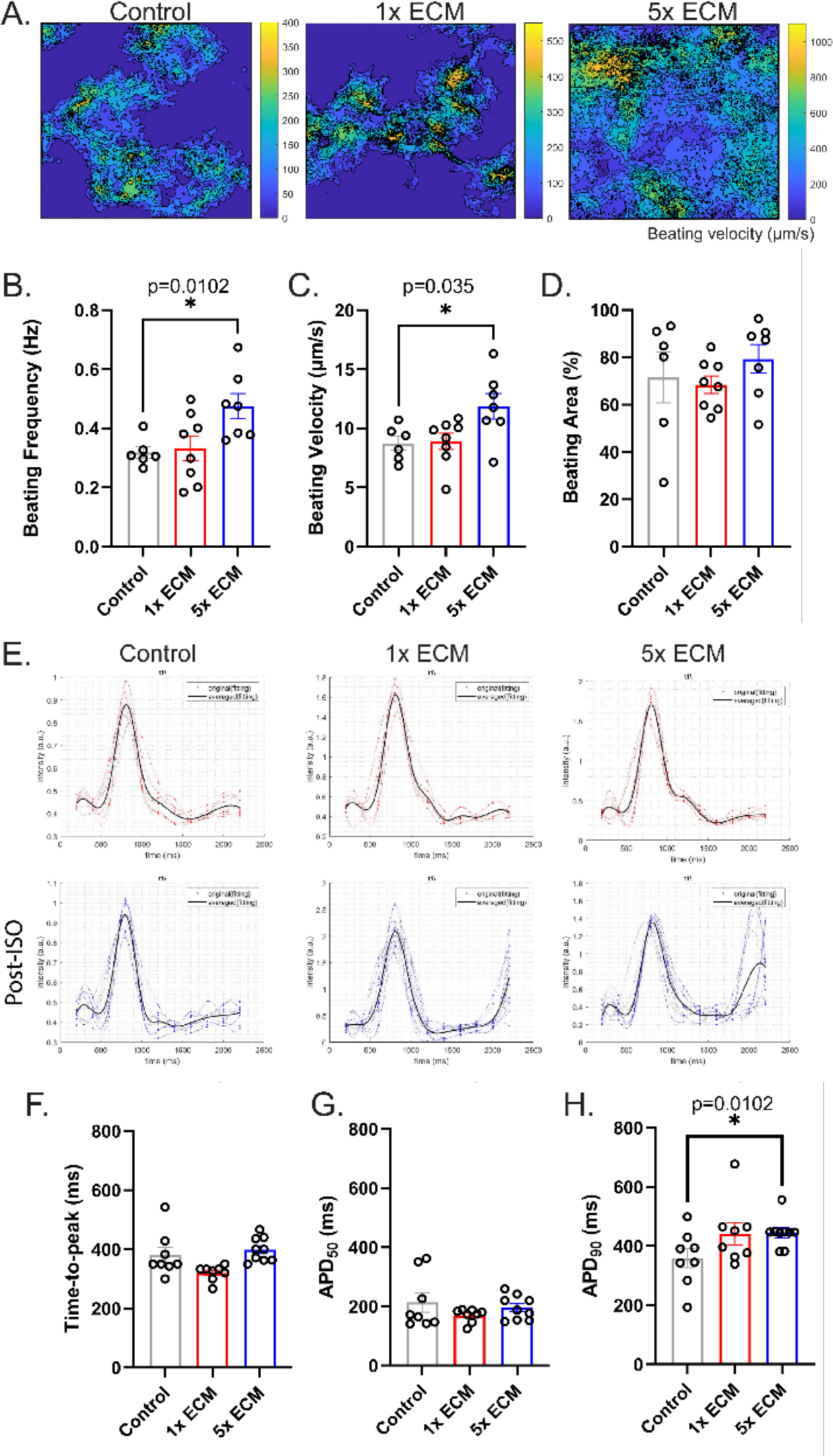
Mechanical and functional assessment of iCMs. (A) Heatmaps showing the beating velocity magnitude and distribution. Quantification of (B) beating frequency, (C) average beating velocity, and (D) beating area. (E) Action-potential curves of iCMs before (red) and after (post-ISO, blue) isoproterenol treatment. Quantification of (F) time-to-peak, (C) 50% decay, and (D) 90% decay. Statistical analysis was done using one-way ANOVA with post-hoc Tukey’s test, *p<0.05, n≥3.

We assessed stimuli responsiveness via changes observed in the beating profile upon isoproterenol (ISO) treatment. Peak-to-peak interval reduced, and the beat frequency increased gradually with ECM pretreatment (Fig. 2E). Collectively these results suggest that high dose ECM pretreatment had a positive impact on the beating kinetics of cells, as well as their responsiveness to stimuli.

### 3.3. Adult ECM pretreatment improves mitochondrial maturity

The contractile function is highly energy dependent. Therefore, we assessed mitochondrial morphology and network structure in iCMs to have an insight into the potential mechanisms underlying the increased function. The control group had significantly low mitochondrial coverage and mitochondria appeared as small spheres or fragmented (Fig. 3A-C). ECM pretreated cells had significantly increased mitochondrial coverage (Fig. 3D). Mitochondria of both ECM pretreated groups appeared elongated, tubular, or filamentous (Fig. 3C). When quantified, high dose ECM pretreated cells had significantly more junction points, hence branching, which indicated more developed, interconnected mitochondrial networks (Fig. 3E-G). These findings suggest that ECM pretreatment increased mitochondrial coverage and complexity in mitochondrial networks. A developed mitochondrial network allows effective distribution of energy substrates and metabolites, suggesting that ECM pretreatment would also enhance the energy metabolism of iCMs.

**Figure 3.**
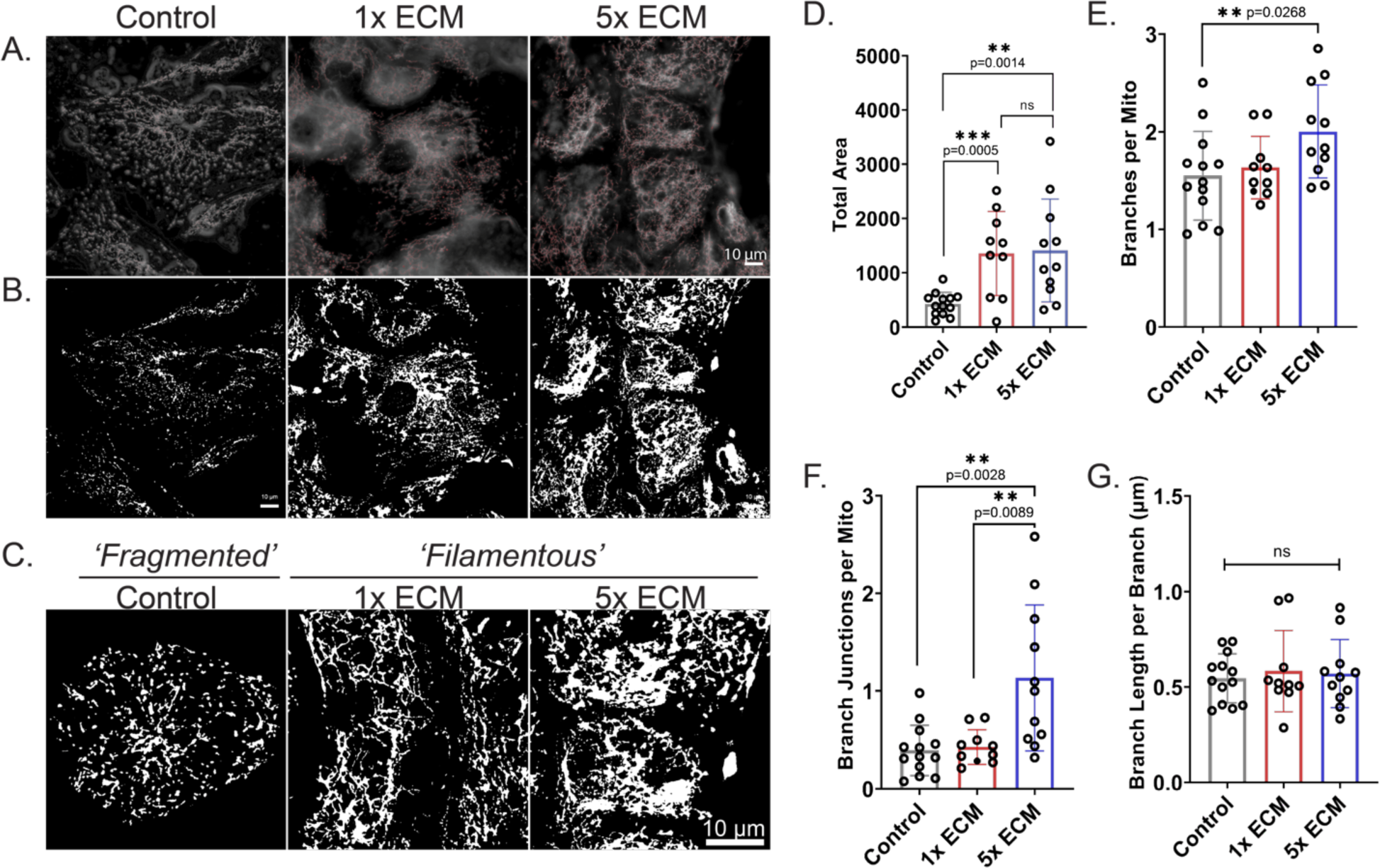
Mitochondrial morphology and network of iCMs. (A) Fluorescence microscopy images of iCMs. Mitochondria were labeled with MitoTracker and imaged with a Zeiss LSM800 laser-scanning confocal microscope and a 1.40-NA 63× objective. Network structure from mitochondrial images superimposed with the mitochondria skeleton (Red) obtained from the Mitochondria Analyzer plugin in ImageJ. (B) Skeleton images of the binary images after thresholding and noise removal (C) Zoomed-in images showing mitochondrial structure. Scale bars represent 10 μm. Quantification of (D) average mitochondrial area in a cell, (E) number of branches of each mitochondrion (F) number of branch junctions of each mitochondrion, (G) the average length of branches of each mitochondrion per branch. Statistical analysis was done using one-way ANOVA with post-hoc Tukey’s test, *p<0.05, **p<0.01, ***p<0.001. n≥3.

### 3.4. ECM pretreatment shifts the energy state

ECM pretreatment generated a more energetic phenotype compared to the control, represented as a dose dependent gradual shift on the energy map (Fig. 4A). Energetic phenotype indicates the utilization of both aerobic and glycolytic metabolic pathways for energy, hence a more mature cardiac metabolism. ECAR values of both glycolysis and non-glycolytic processes (i.e., TCA cycle and glycogenolysis) were increased with high dose ECM pretreatment (Fig. 4B-D). Especially, glycolytic reserve, which indicates the capability of a cell to respond to an energetic demand (Fig. 4C), and non-glycolytic acidification caused by processes in the cell other than glycolysis increased statistically in high dose ECM group (Fig. 4D). ATP production rate also revealed greater ATP production as well as oxidative phosphorylation rate when cells were pretreated with high dose ECM (Fig. 4E). Additionally, gradually increased OCR values of basal and non-mitochondrial respiration were observed in ECM pretreated iCMs (Fig. 4F-I). The energetic demand of the cells significantly increased (Fig. 4G). However, the rate of other cellular oxidative reactions not linked to energy metabolism also increased in ECM pretreatment groups (Fig. 4I). Overall, these results suggest that the high dose ECM pretreatment enhanced the metabolic profile of iCMs by facilitating an efficient utilization of energy substrates.

**Figure 4.**
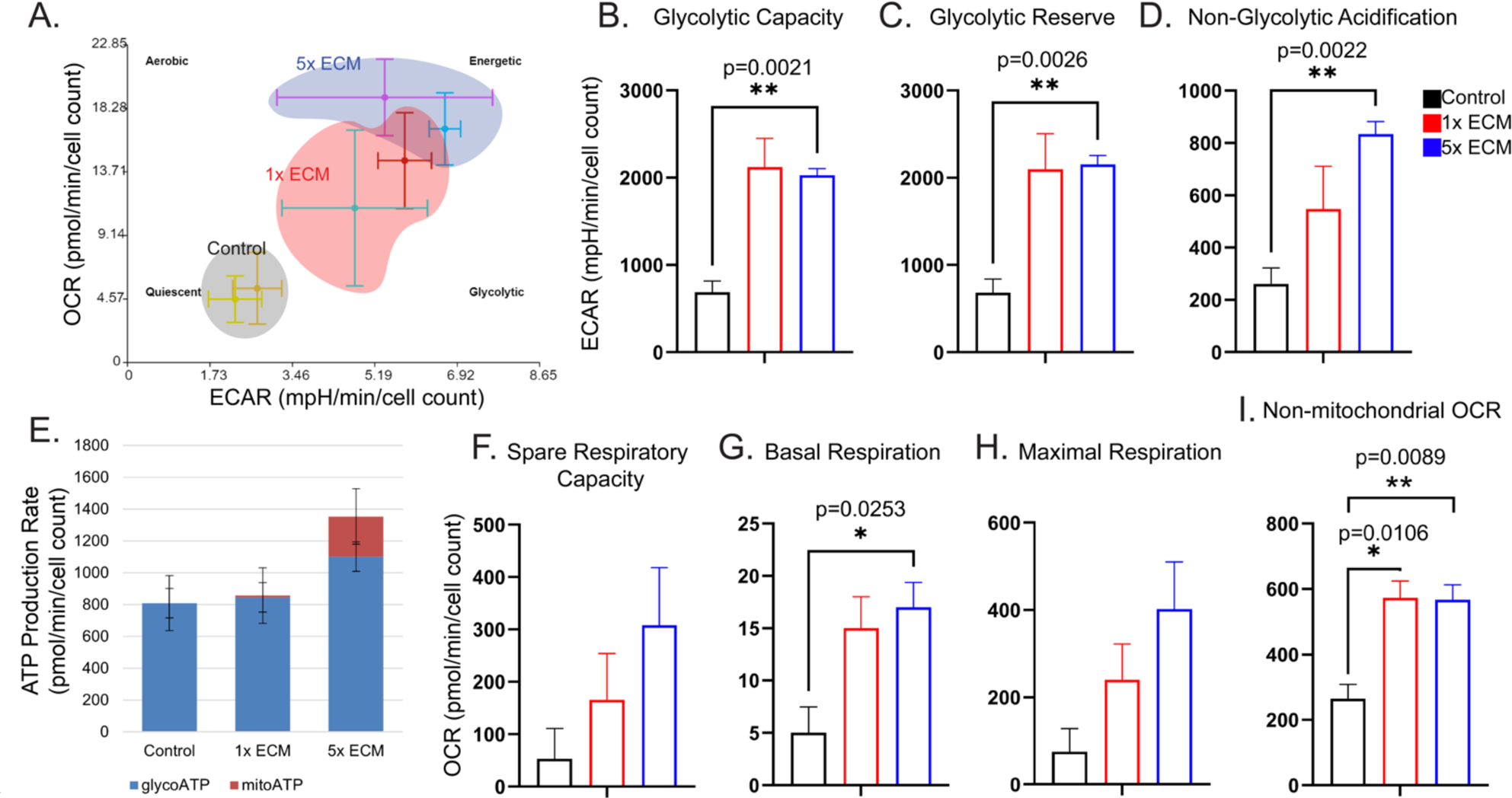
Metabolic profiles of control and ECM pretreated iPSC-CMs. (A) Energy map representing the relative bioenergetic state of the group related to each other. ECAR used for (B) glycolytic capacity, (C) glycolytic reserve and (D) non-glycolytic acidification. E) ATP production associated with the conversion of glucose to lactate (glycolysis, blue) and oxidative phosphorylation (mitochondrial, red). OCR used for (F) spare respiratory capacity, (G) basal respiration, (H) maximal respiration and (I) non-mitochondrial oxygen consumption. n=4 pooled into 2 technical replicates. Graphs constructed with values +-SEM after background correction and cell count normalization.

### 3.5. Gene and protein level characterization

We evaluated the expression levels of a customized panel of human cardiac genes using the NanoString nCounter® technology platform. Applying PCA to this dataset verified the moderate variation (PC1=55%) between ECM pretreated groups and the control group (Fig. 5A). ECM pretreatment lowered expressions of cell proliferation gene *CTGF* and fetal genes including *NKX2-5, ACTA1, ACTA2*, *NPPA* and *NPPB* (Fig. 5B). ECM effect on proliferation was observed in iPSC growth, where high dose ECM treated iPSCs had delayed full confluency compared to the rest (Supp. Fig. 2). Accordingly, gene ontology (GO) analysis of the downregulated genes with ECM pretreatment mapped to canonical Wnt signaling and ECM regulation (Fig. 5C). Wnt signaling inhibition is observed during the cardiac specification after mesoderm formation. Additionally, the downregulation of ECM organization and regulation processes in the pretreated groups suggested the influence of ECM bioactive molecules in the cell media. Moreover, the downregulation of apoptotic signaling pathways emphasized the protective effects of cardiac ECM on the cells.

**Figure 5.**
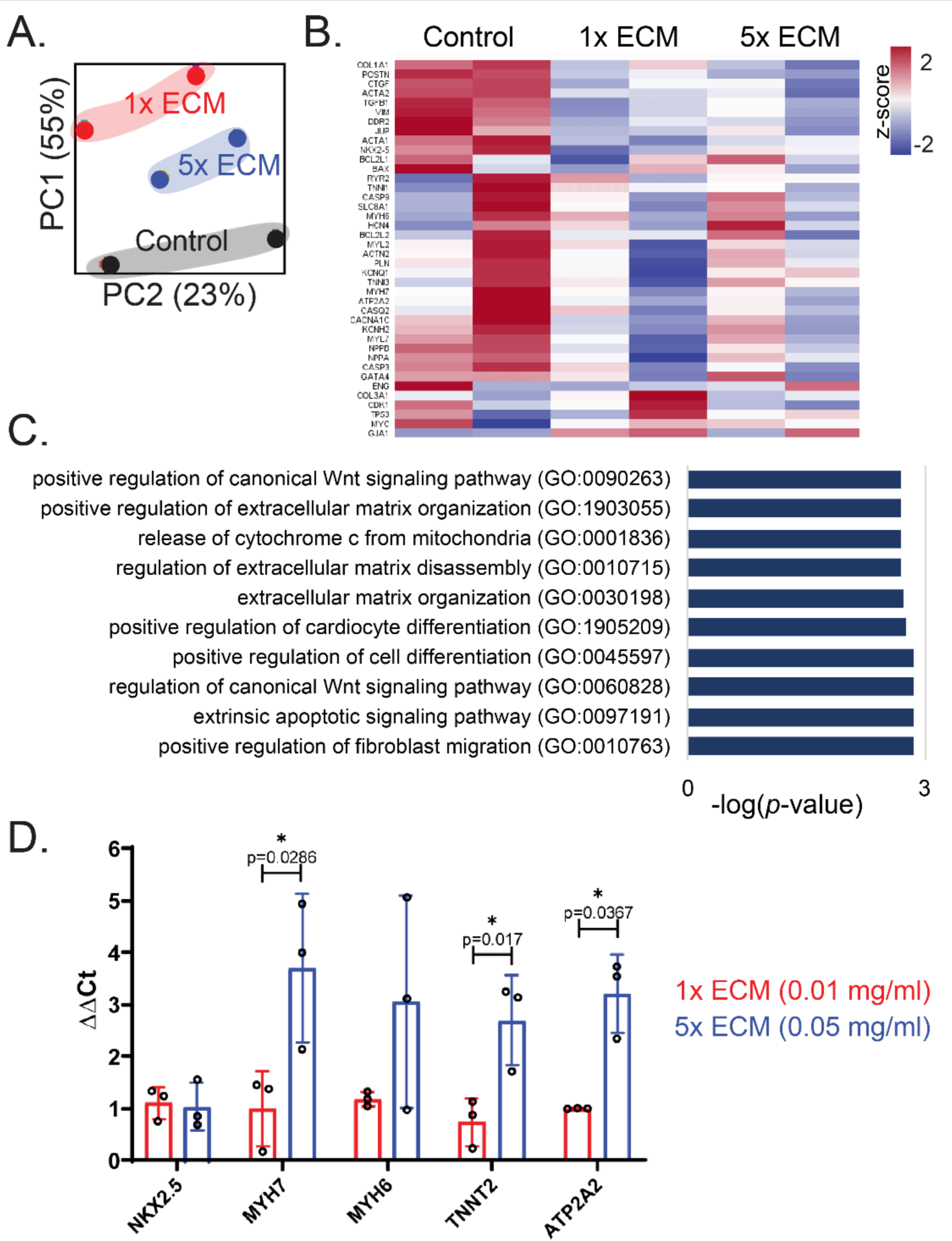
Gene expression up-/downregulated in ECM pretreated iPSC-CMs. (A) Principal component analysis (PCA) on gene expression profiles. The proportion of component variance is indicated as percentages. (B) Heat map of the normalized data. Z-score for gene expression calculated using Euclidean distance and average linkage method. n=6 pooled into 2 technical replicates (C) Top 10 GO biological processes associated with downregulated genes with ECM pretreatment. (D) PCR results for the selected cardiac markers. Data are presented as the mean ± standard deviation. Statistical analysis was done using one-way ANOVA with post-hoc Tukey’s test. *p<0.05, n≥3.

qPCR was performed for a subset of the target mRNA expressions, highly related to cardiac structural and functional maturity. Compared to the control iCMs, high dose ECM pretreated group had upregulated structural MYH7 (p=0.029), TNNT2 (p=0.017) and functional ATP2A2 (p=0.037) expressions (Fig. 5D). These results indicate that ECM pretreatment might affect cardiac differentiation dynamics, as well as structural and functional maturity of resulting iCMs.

### 3.6. Investigation of the ECM components responsible for the observed effects

The ECM contains all four major classes of biological macromolecules, with proteins being the most abundant. To investigate whether the observed effects were caused by the protein component, we subjected the ECM to heat denaturation. Additionally, we sonicated ECM to release the contents of extracellular vesicles (EV) that might be bound within the matrix, thereby enhancing their accessibility to cells. Moreover, as a control for the sonication group, we added EVs derived from human LV. As a secondary measure, we followed an alternative decellularization and digestion protocol, which aligns with current standards for preserving the EV integrity (Peracetic acid, PAA groups).

**Supplementary Figure 2.**
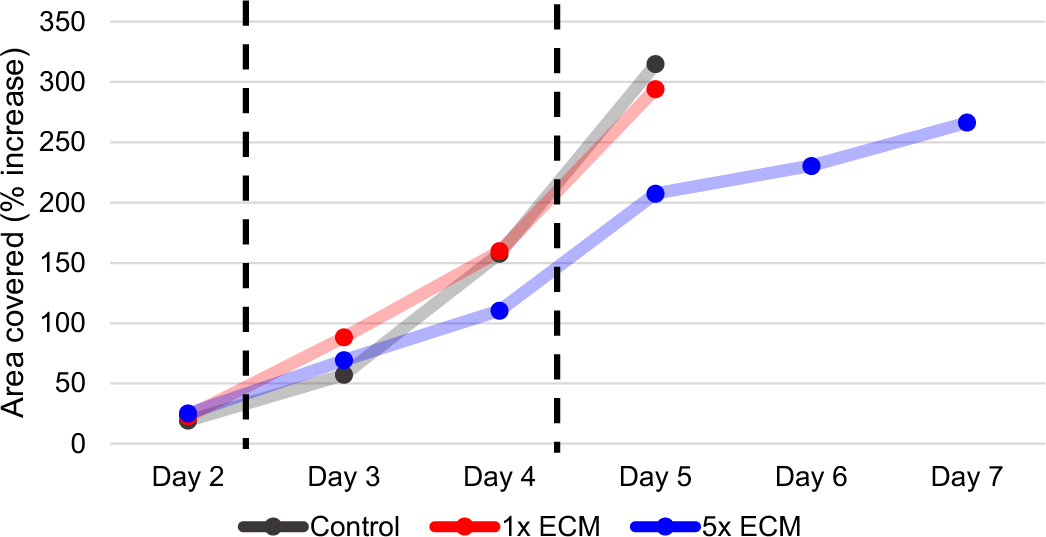
Increase in area covered by iPSCs until they reach confluency. All data were normalized to the initial area coverage.

Beating frequency showed an increasing trend across the groups, from the untreated control to the heat-denatured ECM, ECM, sonicated ECM, and ultimately the EV (Fig. 6A). However, no significance in beating kinetics was detected within groups. At the subcellular level, mitochondrial coverage was also comparable across the groups, yet ECM, sonicated ECM and EV pretreated iCMs had more developed mitochondrial networks. Branching and interconnectedness significantly increased in these groups (Fig. 6B-C). However mitochondrial development didn’t translate into the energy profile of these cells. Only the EV treated group was separated from the rest, showing higher variability spanning from aerobic to energetic profile (Fig. 6D and Supp. Fig. 3). We did not record any noticeable benefit of the PAA decellularization method over the standard ECM decellularization method used in the previous sections.

**Figure 6.**
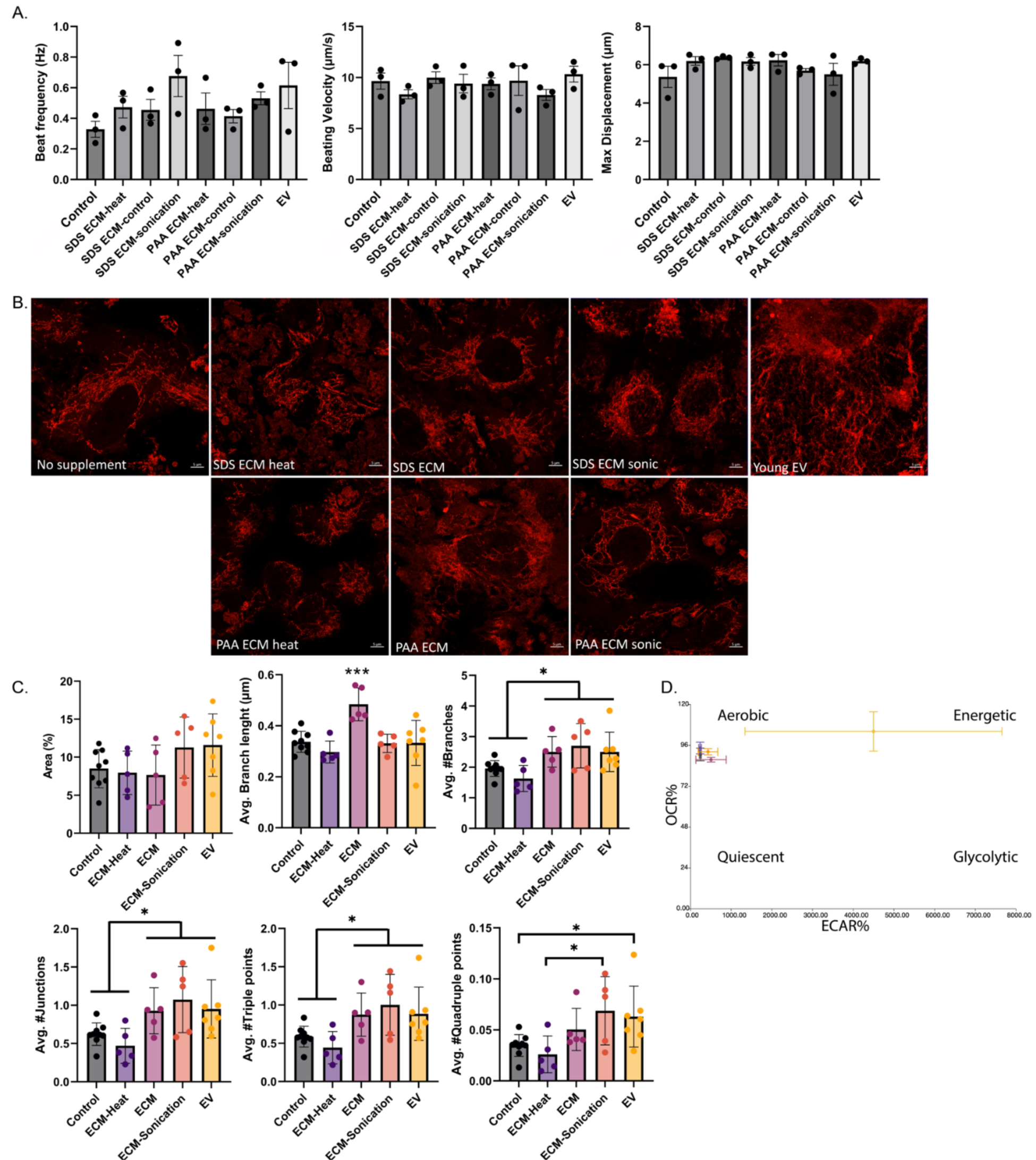
Quantification of (A) beating frequency, beating velocity and maximum displacement of spontaneous beating. Control is no ECM pretreated group and all ECM groups are high-dose (5X). (B) Mitochondrial morphology and network of iCMs. (A) Fluorescence microscopy images of iCMs. Mitochondria were labeled with MitoTracker and presented as skeleton images of the binary images after thresholding and noise removal. Scale bars represent 5 μm. Quantification of (C) average mitochondrial area in a cell, the average length of branches of each mitochondrion per branch, number of branches, junctions, triple and quadruple points of each mitochondrion. Statistical analysis was done using one-way ANOVA with post-hoc Tukey’s test, *p<0.05, **p<0.01, ***p<0.001. n≥3. (D) Energy map representing the relative bioenergetic state of the groups.

**Supplementary Figure 3.**
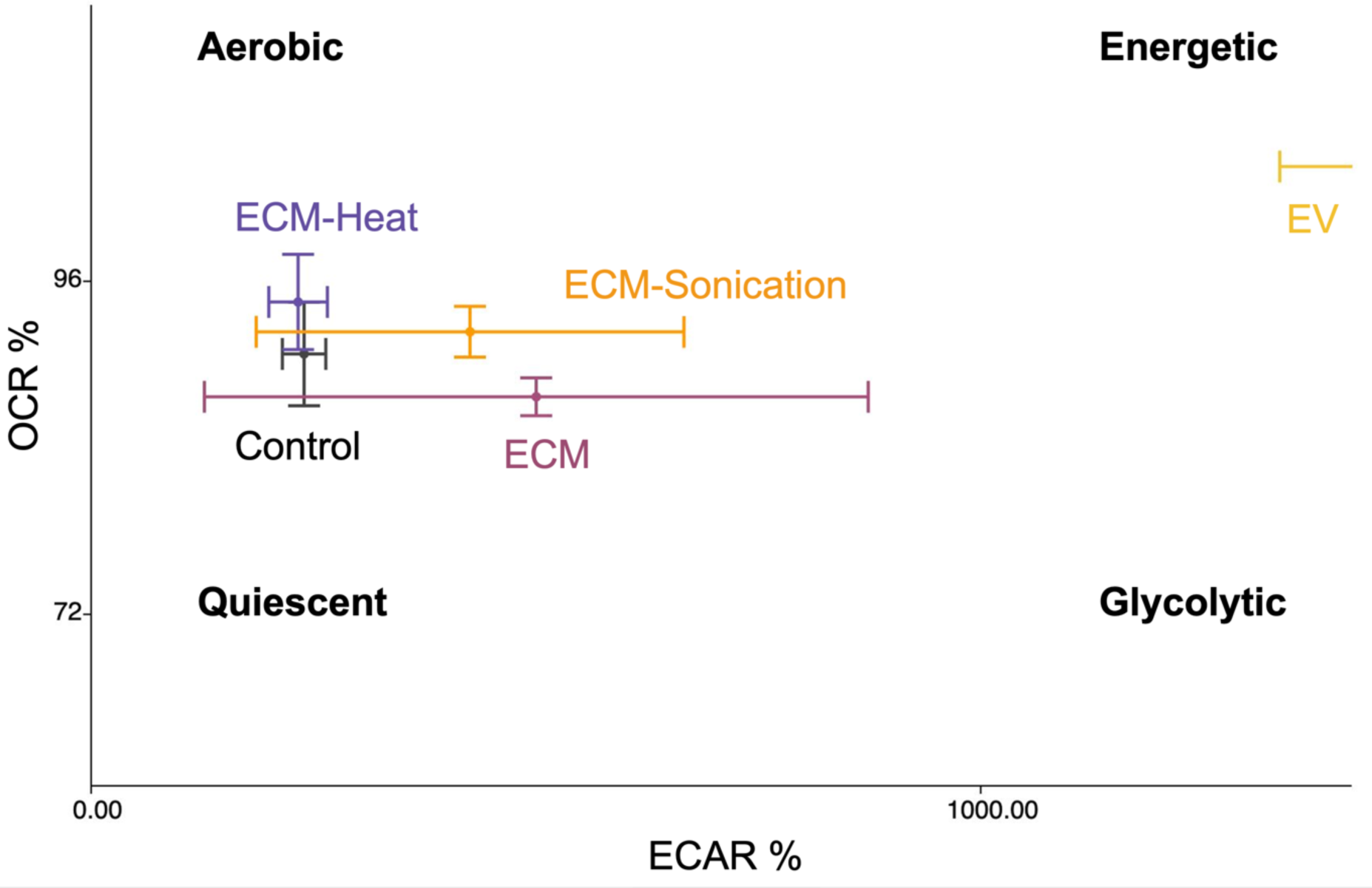
Zoomed in image from the energy map, showing the relative bioenergetic state of the groups (Control=black, heat-denatured ECM=purple, ECM=pink, sonicated ECM=orange, EV=yellow).

## 4. Discussion

Human iCMs provide an attractive cell source for heart regeneration, along with disease modeling, drug screening, and toxicity testing. Despite an extensive effort to enhance cardiac maturity[9,29], the achieved CM maturity has been limited mainly to structural aspects, leaving functional and metabolic maturation as attractive targets. Here, we conditioned iPSCs with adult human heart derived ECM at the pre-differentiation stage to generate functionally and metabolically mature iCMs. The resulting cells exhibited increased beating kinetics, drug responsiveness (Fig. 2), mitochondrial coverage and network complexity (Fig. 3) as well as a metabolic shift toward an energetic state (Fig. 5). Additionally, ECM pretreated iCMs exhibited increased expression of mature structural and functional markers while reducing cell proliferation and fetal cardiac markers (Fig. 4). Moreover, our results provided additional evidence to suggest that the observed effects were not solely due to the ECM proteins (Fig. 6). Instead, other ECM components, which are less or non-susceptible to denaturation (i.e., HSPGs) may play more significant roles in affecting cell fate determination, as recently reported by others[30]. Our findings are a step forward in enhancing cardiac maturation not just functionally but also energetically. Here, we provided a rather easy and rapid protocol to follow to overcome the current CM immaturity limitations.

Decellularized ECM has been used to mimic the natural microenvironment and to study cell-ECM interactions [31–33]. Relatedly, in a recent study, ECM from adult and aged tissues (39-84-year-old for all three tissues, 52-70-year-old for heart tissues only) was reported to carry biochemical factors that direct tissue-specific differentiation in multiple organs [30]. However, note that our previous findings showed that cell-ECM interplay is highly dynamic and age-dependent[21,34,35]. A comparison between young (<30-year-old) and aged (>50-year-old) human hearts revealed decreased glycoproteome in aged tissue, which was linked to age-related functional decline in a mouse model [36]. Moreover, we found that accumulated toxic waste lipofuscin was retained in aged tissue even after decellularization [35]. This is especially important that lipofuscin, being rich in heavy metals, causes oxidative damage to cells, impairs cardiac and mitochondrial function and induces cell death [37,38]. Additionally, iCMs cultured on aged ECM exhibited elongated sarcomeres comparable to native adult CMs but also aging and stress markers [2]. Thus, grouping adult and aged ECMs could lead to confounding results, and aged ECM may not be suitable for long term cell conditioning or treatment.

Adult ECM, on the other hand, has shown promise in promoting iCM structure, maturity and function [2,39,40] and supporting iPSC and ESC differentiation in various systems [30,41–45]. Notably, although these studies do not explicitly label the ECM as adult due to the lack of other age group comparisons, the majority utilized ECM derived from adult tissues (i.e., 3-6-month-old pig) due to convenience and ease of access. While ECM is known to affect iPSC differentiation, it is still poorly understood *how* it specifically supports CM differentiation. Previous studies reporting enhanced cardiac differentiation or maturation focus mainly on electrophysiological function, Ca2+ handling abilities and cardiac marker expression. Here, we used ECM derived from adult human hearts (30-50-year-old) to condition iPSCs and evaluated iCM maturity at multi-levels, including sarcomeric structure, beating and calcium handling abilities, transcription factor activation, as well as bioenergetics and metabolism, which have recently gained significant attention in the field. Investigating the impact of ECM pretreatment on the mitochondria and metabolic transitions of iCMs provided a novel aspect that elucidates the underlying reasons for the observed functional improvements by the adult ECM.

Higher TNNT2 expression suggested that ECM pretreated iCMs achieved a higher degree of cardiomyocyte differentiation and have progressed further along the maturation pathway, resembling more mature and functional cardiac muscle cells (Fig. 5). Additionally, high expression of MYH7, isoform with higher ATPase activity, and ATP2A2 suggested an increased contractile efficiency and functional maturation (Fig. 2 and 5D). CMs that expressed adult type MYH isoform was previously shown to demonstrate greater shortening velocities and generated more peak power[46]. Similarly, we observed increased beating/shortening velocity (Fig. 2C) and greater peak calcium intensity (Fig. 2E) in ECM pretreated iCMs, indicative of improved contractile properties. Moreover, faster Ca2+ uptake facilitated by SERCA2 (coded by ATP2A2) contributed to efficient repolarization of iCMs as well as drug sensitivity (Fig. 2F). APD90 was prolonged as iCMs matured in the presence of increasing cytosolic Ca2+ activity (Fig. 2H) and a ventricle-type AP peak with a shoulder appeared (Fig. 2E).

The enhancement in contractile properties and functional maturation is closely tied to bioenergetic and metabolic adaptations. These interdependent processes work synergistically to meet the increased energy demands during cardiomyocyte differentiation and maturation *in vivo*. Notably, here we showed that ECM pretreated iCMs had increased mitochondrial coverage (Fig. 3A-D) and improved mitochondrial network complexity (Fig. 3E-G). Elongated, interconnected mitochondrial networks suggested active mitochondrial fusion, which is advantageous for efficient energy production through oxidative phosphorylation and calcium buffering [47,48]. Building on previous research showing decreased glucose uptake and increased fatty acid consumption in iPSCs when differentiated in the presence of pig ECM using colorimetric assays [49], we conducted a thorough analysis of the energy profiles of iCMs pretreated with human ECM. Our findings revealed an ECM-dose dependent gradual metabolic shift towards an energetic state (Fig. 4A). Increased glycolytic capacity and reserve mimicked early cardiac development, implying an increased energy demand from glycolysis to support CM proliferation and differentiation (Fig. 4B-C). Relatedly, increased non-glycolytic acidification can be attributed to enhanced oxidative phosphorylation or cellular processes such as protein synthesis (Fig. 4D-E). Moreover, the gradual increase in the spare respiratory capacity as well as basal and maximal respiration suggested that ECM pretreated iCMs were at a dynamic and adaptive metabolic state, providing higher flexibility to accommodate increasing energy demands (Fig. 4F-H). However, we acknowledge that adult ECM pretreatment did not generate fully mature iCMs, and the non-mitochondrial oxygen consumption results support this. An increased non-mitochondrial OCR combined with increased glycolytic and mitochondrial ATP generation suggests that iCMs were most likely in transition and that their mitochondrial respiration was not fully established yet (Fig. 4I). Overall, we outlined the intricate relationship between enhanced contractile properties, calcium handling and the associated transcriptomic, mitochondrial and metabolic adaptations that occurred upon ECM pretreatment during pre-differentiation stage. Altogether these findings confirm the robustness of adult ECM treatment approach to improve cardiac differentiation and subsequent maturation.

Regarding *what* causes these observed effects, there are extensive studies investigating and comparing ECM substrates (i.e., Laminin-521, -221, Collagen-I, Collagen-IV, Fibronectin, Vitronectin) alone or in combination, for their ability to support iCM differentiation[50]. Certain ECM proteins, such as fibronectin, have been reported to promote the formation of precardiac mesoderm, while others, like laminins (Laminin-111, -521) and collagen-IV, have been found to inhibit cardiogenesis [51]. However, these studies typically focus on early cardiac differentiation (i.e., analysis on 15-day-old iCMs) hence CM maturity was often not reported. Note that studies focus on identifying the key ECM *protein* or its derivatives to drive the iCM differentiation. However, proteins are not the only ECM components. To isolate the effect of ECM proteins including ECM cytokine and chemokines, we heat denatured ECM (ECM-Heat) and treated iPSCs with this heat denatured ECM (Fig. 6).

Most proteins undergo denaturation at temperatures exceeding their physiological range (> 60°C). However, the rate at which collagen fibrils denature was reported to depend not only on temperature and time but also on the environment being wet (i.e., water bath) or dry (i.e., oven). It was shown that collagen fibrils can maintain their native structure after being heated to 120°C for 90 min in the absence of water, whereas no collagen fibers were left when heated to 80°C for 30 min in a wet environment [22]. The impact of heat extends to other ECM components that contain carbohydrates (i.e., GAG) such as proteoglycans and glycoproteins. Despite their relatively higher heat stability, heat treatment can still lead to the fragmentation of these components into smaller molecular weight fragments [52], potentially altering their function and interactions. To selectively mitigate the influence of heat-sensitive ECM components, mainly ECM proteins, we subjected ECM to a 3-hour heat treatment at 80°C in a water bath.

The energy state of the untreated iCMs (Control) was comparable to the heat treated ECM (ECM-Heat) group (Fig. 6D) and at a lower energy state than the rest (Supp. Fig. 3). Relatedly, although heat treatment of ECM didn’t negatively affect the beating properties of ECM treated cells, we noted a significant drop in the mitochondrial network complexity (Fig. 6B-C). ECM pretreatment increased mitochondrial network complexity and interconnectivity, however heat-denaturation eliminated the increased mitochondrial branching effect seen with ECM treatment, yielding similar results to those of the Control group (Fig. 6C). These results suggest that collagens, which are the most abundant proteins in the ECM, and other heat sensitive ECM components including growth factors may have a reduced role in the overall cardiac functional while affecting mitochondrial structure by either promoting mitochondrial fusion or inhibiting mitochondrial fission. Growth factors found in the human heart ECM, namely FGF19, TGFb and PDGF, [35], were shown to increase mitochondrial biogenesis and fusion in various cells [53–55]. Alternatively, while the exact connection between ECM structural proteins (i.e., collagen-I, fibronectin and vitronectin) and cardiolipin is not completely understood, cardiolipin, a mitochondria-exclusive phospholipid responsible for both mitochondria fusion and fission [56], was reported to inhibit cell attachment to these ECM proteins[57]. This suggests that the presence of ECM proteins might impact mitochondrial network structure through cardiolipin. However, further research is needed to elucidate the nature of this relationship or whether any such connection exists. Nonetheless, these findings indicated that the abundant ECM proteins are not the sole effectors, and the beating and energy metabolism were minimally affected (Fig. 6 and Supp. Fig. 3) when proteins were denatured.

The beneficial effects of ECM were suggested to be conveyed by the extracellular vesicles (EV)[58]. Therefore, we investigated whether we could amplify the observed effects by increasing the availability of ECM-bound vesicles’ cargo (i.e., miRNA, growth factors) to the cells. To release ECM-bound factors and disrupt the EV membrane integrity [59], we sonicated ECM on ice (ECM-Sonication). In a prior study, we verified the preservation of ECM-bound exosome-like EVs in decellularized cardiac tissues [21]. However, due to the challenges in EV isolation and harvesting, we followed two decellularization methods: ionic/nonionic detergent decellularization followed by trypsin digestion (SDS-ECM) and peracetic acid/ethanol decellularization followed by collagenase digestion (PAA-ECM) (Fig. 6). Doing so ensured that the methods used for tissue handling did not limit the amount and quality of EVs in our study.

Pretreatment with ECM-Sonication resulted a similar effect on cells as isolated EVsAlthough not significant, beating frequency showed an increasing trend towards ECM-Sonication and EV (Fig. 6A) and both groups showed an increase in mitochondrial area coverage and complexity (Fig. 6B-C). These changes indicated an increased energy adaptation, however, only EV pretreated cells exhibited a wide-spanning metabolic profile towards an energetic state (Fig. 6D). Overall, these results suggest that the factors tightly bound to the ECM (i.e., growth factors) are preserved and released after sonication to perform similarly to isolated EVs. Moreover, EV cargo is beneficial but not the dominant force driving cardiac fate and subsequent cardiac maturation.

We identified high amounts of collagen-I and collagen-IV, along with carbohydrate- and lipid-binding proteins in adult ECM (Fig. 1D) that have direct or indirect effects in promoting the initial stages of iPSC cardiac differentiation and/or cardiac metabolic maturation. For instance, fibronectin was associated with ILK signaling cascades and IGF-1 transport and uptake (Supp. Fig. 1B), which in turn promotes mesoderm formation for an efficient cardiac differentiation [51] and boosts mitochondrial function hence oxidative metabolism during cardiomyocyte adaptive growth by a Ca2+ uptake-dependent mechanism in cultured human and rat CMs [60], respectively. Similarly, apolipoproteins were associated with enhanced cardiac differentiation of ESCs and iPSCs and promoted maturation of the Ca2+ handling properties of ESC-derived CMs via the BMP4/SMAD signaling pathway [61]. Moreover, HSPGs and glycan-binding lectins (i.e., Galectin) were mainly associated with cardiogenesis. Perlecan and Glypican-3 deficiency was reported to result in defective heart development in mice[28]. Another basement-membrane-specific HSPG, agrin was also identified as a determinant of normal heart development and crucial regulator of epicardial epithelial-to-mesenchymal transition (EMT) in both human and mouse models[62]. Further research is needed to dissect the individual effects of ECM components, however, these identified components seem to affect cardiac differentiation and maturation via various metabolic pathways. Referring to the heat denatured ECM results, the observed effects might be related not to the protein core but to the decorative units attach to it.

While the study focused on the biochemical aspects of ECM, it is essential to acknowledge that ECM biophysical properties, such as stiffness, tensile strength, architecture and orientation also play a crucial role in cell-ECM dynamics and stem cell differentiation. Solubilizing ECM sacrificed these valuable effects of intact tissues. Nonetheless, the fact that we could still capture the beneficial effects of adult heart derived ECM on cardiac differentiation and enhancing iCM maturity is impressive. It is evident that early exposure to adult ECM drives iPSCs for a tissue-specific fate and yields functionally and metabolically more mature CMs. Moreover, this effect interestingly seems to be independent of the source (human vs xenogeneic), form (solution vs powder) and induction timing (before or during differentiation)[30,49].

Finally, a direct comparison of ECM from other tissues or xenogeneic sources was not performed in this study. However, advancements in using xenogeneic organs/scaffolds in tissue engineering[63–66], and reported high similarity between human and pig heart matrix compositions[49] indicate the potential of using xenogeneic cardiac tissue for commercial iCM maturation purposes. Further research is still needed to investigate potential interspecies differences in tissue signatures and long-term effects of xeno-ECM. Nonetheless, this collective consensus on the early use of adult ECM to enhance tissue specialization of iPSCs is very promising. This novel protocol could significantly minimize the time and money spent on cardiac maturation protocols and put us a step forward in repopulating post-MI hearts with patients’ own cells.

## Conclusion

In conclusion, our study offers adult human heart-derived ECM pretreatment as an easy-to-follow protocol to direct iPSCs toward a cardiac fate and enhance cardiac maturation. This protocol holds great promise in reducing the time and resources required for cardiac tissue engineering and regenerative therapies, bringing us closer to the prospect of repopulating post-myocardial infarction hearts with patient-specific cells. Moreover, the insights gained from this study have broader implications for in vitro cardiac disease modeling and drug screening studies. By using these functionally and metabolically mature iCMs, more reliable and clinically relevant results can be obtained in vitro. Further research into specific molecular mechanisms underlying the effects of ECM pretreatment will be crucial to fully unlock the potential of this novel protocol.

## Author contributions

S.G.O. and P.Z. designed research, S.G.O. performed research, S.G.O. and M.T. analyzed data, S.G.O., M.T. and P.Z. conducted review and editing, P.Z. provided funding, project administration, and resources, S.G.O. wrote the paper and P.Z. revised the paper.

## Ethics Statement

Deidentified human hearts were collected through the Indiana Donor Network under the Institutional Review Board (IRB) approval for deceased donor tissue recovery. All human tissue collection conformed to the Declaration of Helsinki.

## Data Availability Statement

The raw/processed data required to reproduce these findings can be shared upon reasonable request.

## Acknowledgment

This work was funded by the NSF-CAREER Award No 1651385 and NIH Award 1 R01 HL141909-01A1. We thank Indiana Donor Network and Dr. Keith L March for the human heart tissues.

---

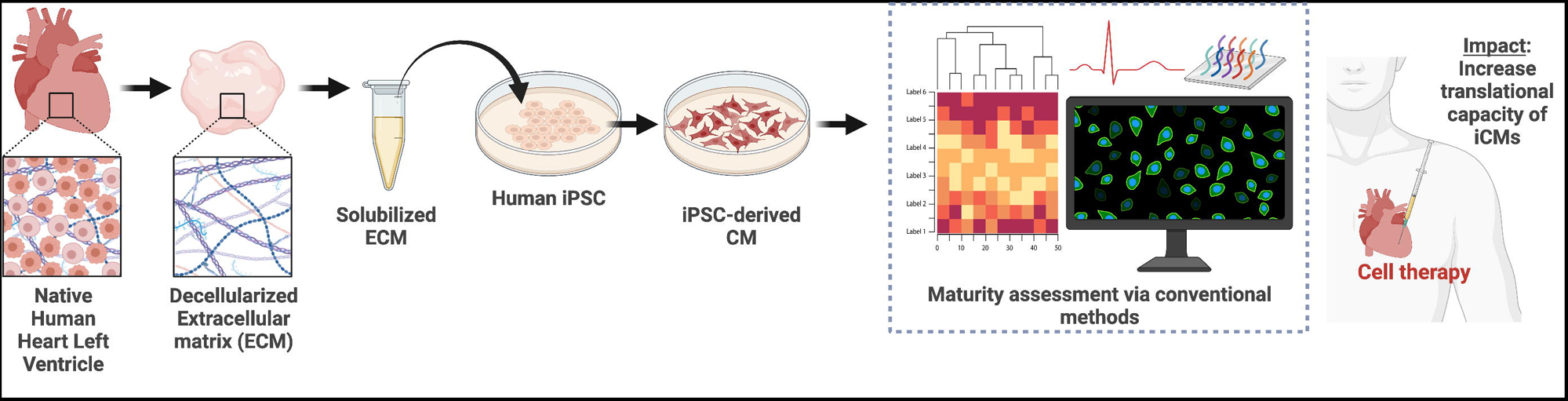

